# GeneFíor: A back to basics and transparent multi-tool approach to sequence detection

**DOI:** 10.64898/2026.05.15.724838

**Authors:** Nicholas J. Dimonaco, Katie Lawther

**Affiliations:** Institute for Global Food Security, School of Biological Sciences, Queen’s University Belfast, Belfast, UK; Department of Computer Science, Aberystwyth University, Aberystwyth, UK; Laboratory of Microbiology, Wageningen University, Wageningen, The Netherlands

## Abstract

The detection of sequences of interest, such as antimicrobial resistance genes, directly from genomic and metagenomic sequencing data has become routine, enabled by curated reference databases and rapid *in silico* sequence search tools. Yet most workflows depend on prior assembly, an inherently lossy process in which a substantial proportion of reads fail to assemble or are collapsed into consensus sequences, causing low-abundance variants and nucleotide-level diversity to be systematically obscured. The tools used to interrogate the resulting assemblies compound this further, clustering reference sequences at arbitrary identity thresholds, imposing hidden parameter defaults, and reducing intermediate alignment evidence to summarised outputs that cannot be critically evaluated or reproduced.

Here we present GeneFíor, a transparent, multitool workflow integrating BLAST, DIAMOND, Bowtie2, BWA, and Minimap2 to search both DNA and protein sequences against any user-supplied reference database. By enforcing genecentric identity and coverage thresholds at both the read and gene level, GeneFíor reduces false positives while retaining sensitivity to genuine, low-abundance variants, including those differing at single-nucleotide resolution. Crucially, by exposing all alignment parameters, preserving intermediate outputs, and generating cross-tool consensus detection matrices, GeneFíor makes the influence of tool choice, database selection, and parameter configuration on reported gene profiles directly observable and reproducible.

## 1 Introduction

Gene detection from genomic and metagenomic data underpins a wide range of biological analyses, from clinical antimicrobial resistance (AMR) surveillance to environmental and industrial microbiology. Assembly-based approaches are often treated as a prerequisite for high-confidence detection, yet they are inherently lossy: substantial proportions of reads fail to assemble, while others are collapsed into consensus sequences that mask underlying variation. This is particularly problematic in complex microbial communities, where only a biased subset of the original data is retained. Low-abundance variants, including those differing at single-nucleotide resolution, are routinely lost as a consequence. The ability to identify genes directly from raw reads, without reliance on assembly, is therefore both critical and technically challenging.

Despite the availability of numerous specialised tools for read-based gene detection, issues of transparency, reproducibility, and database flexibility persist. Methods that perform direct read-to-reference mapping frequently rely on hidden parameter defaults, cluster reference sequences using arbitrary thresholds, and reduce intermediate results to summarised outputs. These shortcomings make it difficult to assess, reproduce, or critically evaluate reported gene profiles. As a result, detected gene sets often reflect methodological choices and tool behaviour as much as they do the underlying biology.

These limitations are particularly consequential in the context of antimicrobial resistance gene (ARG) detection, where the stakes of inaccurate or irreproducible results are high [1, 2]. Standardised laboratory methodologies for identifying and classifying resistant organisms are well developed [3, 4, 5], and although not without limitations [6], they provide clearly defined parameters that enable reproducible and interpretable results. In contrast, *in silico* approaches for ARG detection have expanded rapidly, promising not only gene detection from sequencing data [7], but also phenotype prediction from genomic content [8, 9, 10]. Curated databases such as ResFinder [8] and CARD (Comprehensive Antibiotic Resistance Database) [9] have enabled rapid growth of *in silico* ARG profiling, and a diverse tool landscape has emerged, offering proprietary databases [11], mutational marker detection [12], variation graph approaches [13], and machine learning classifiers [7]. However, these approaches often rely on opaque algorithmic choices, reference sequence clustering decisions and reporting practices that obfuscate the confidence of reported hits and are not always clearly communicated to the users. Yet despite this apparent diversity, most tools rely on a common foundation of established search algorithms: BLAST [14], DIAMOND [15], Bowtie2 [16], BWA [17], and Minimap2 [18], whose behaviour is well characterised and whose outputs are comparatively interpretable. This growing preference for opaque, highly summarised reporting sits in tension with this foundation, and obscures biologically meaningful variation: resistance emerges through diverse mechanisms including horizontal gene transfer, chromosomal mutation, and ecological selection [19], and similar phenotypes can arise from fundamentally different genetic events [20], making nucleotide-level transparency essential. This proliferation of increasingly complex and opaque methods raises a fundamental question: in the pursuit of novelty and performance, have we compromised interpretability, reproducibility, and user control?

The same challenges extend beyond AMR. Databases such as HADEG (Hydrocarbon Aerobic Degradation Enzymes and Genes) [21] and VFDB (Virulence Factor Database) [22] support functional profiling relevant to bioremediation and virulence, yet the tools used to interrogate them suffer from identical limitations: inflexible pipelines, poor parameterisation, and restricted database compatibility. Researchers across domains face the same core problem: rigorous and reproducible gene detection remains elusive.

To address this, we present GeneFíor (pronounced Gene-feer; *fíor*, Irish for ‘true’), a transparent and flexible multi-tool workflow for accurate and interpretable gene detection from both DNA and protein sequence data. GeneFíor integrates BLAST, DIAMOND, Bowtie2, BWA, Minimap2, and MMseqs2 within a unified framework, enabling users to interrogate any curated database with full control over alignment parameters and detection thresholds. By enforcing target-gene-centric coverage and identity validation, GeneFíor reduces false positives while retaining sensitivity to low-abundance variants, and all intermediate outputs are preserved to ensure complete analytical transparency. Crucially, all intermediate outputs and mapping statistics are preserved, ensuring complete analytical transparency. Additional modules, including GeneFíor-Recompute and GeneFíor-Gene-Stats, further support detailed interrogation and reproducibility of results. We also provide AMRFíor, a dedicated submodule for ARG detection querying CARD and ResFinder by default, which we provide a use-case here.These results demonstrate how tool, sequence type, and parameter choices measurably shape reported resistomes.

Rather than obscuring methodology behind complex pipelines or machine learning models, GeneFíor provides a back-to-basics and transparent multi-tool consensus detection approach with user-defined parameters and comprehensive output.

## 2 Methods

### 2.1 GeneFíor

GeneFíor is a transparent, modular workflow for gene detection directly from unassembled DNA sequence reads and predicted gene sequences. Rather than introducing a novel alignment algorithm, GeneFíor is designed around the principle that the most reliable and interpretable gene detection is achieved by harnessing well-validated, widely adopted sequence search tools in a unified, user-controlled framework. To this end, GeneFíor integrates five established sequence alignment and search algorithms, BLAST v2.17.0, DIAMOND v2.1.13, Bowtie2 v2.5.4, BWA v0.7.19, and Minimap2 v2.30, enabling simultaneous interrogation of both nucleotide and amino acid sequences against any user-supplied reference database.

A central design objective of GeneFíor is transparency. All intermediate outputs, per-tool alignment statistics, and detection metrics are retained and made accessible to the user, in contrast to tools that obscure these behind summarised or black-box outputs. Detection thresholds, including minimum sequence identity and gene coverage, are fully user-configurable rather than fixed, enabling researchers to tailor sensitivity and specificity to the biological question at hand. Gene detection is governed by a gene-centric coverage validation strategy (Section 2.3), which requires alignments to meet both identity and breadth-of-coverage criteria before a gene is reported as detected, mitigating false positives arising from spurious matches to conserved domains or short homologous regions. GeneFíor generates a cross-tool consensus detection matrix (Section 2.4) that summarises which genes are identified by each algorithm, allowing users to assess concordance across tools and apply multi-tool agreement as an additional confidence filter. This design enables direct comparison of how tool choice, database selection, and parameter settings influence reported gene profiles, a capability that is rarely afforded by existing specialised detection tools.

GeneFíor is written in Python and is available via PyPi (https://pypi.org/project/GeneFior/), Bioconda (https://anaconda.org/channels/bioconda/packages/genefior/overview and GitHub (https://github.com/NickJD/Genefior). Installation is recommended through Bioconda as all search and processing tools are installed together. A high-level overview of GeneFíor is shown in Figure 1.

**Figure 1:**
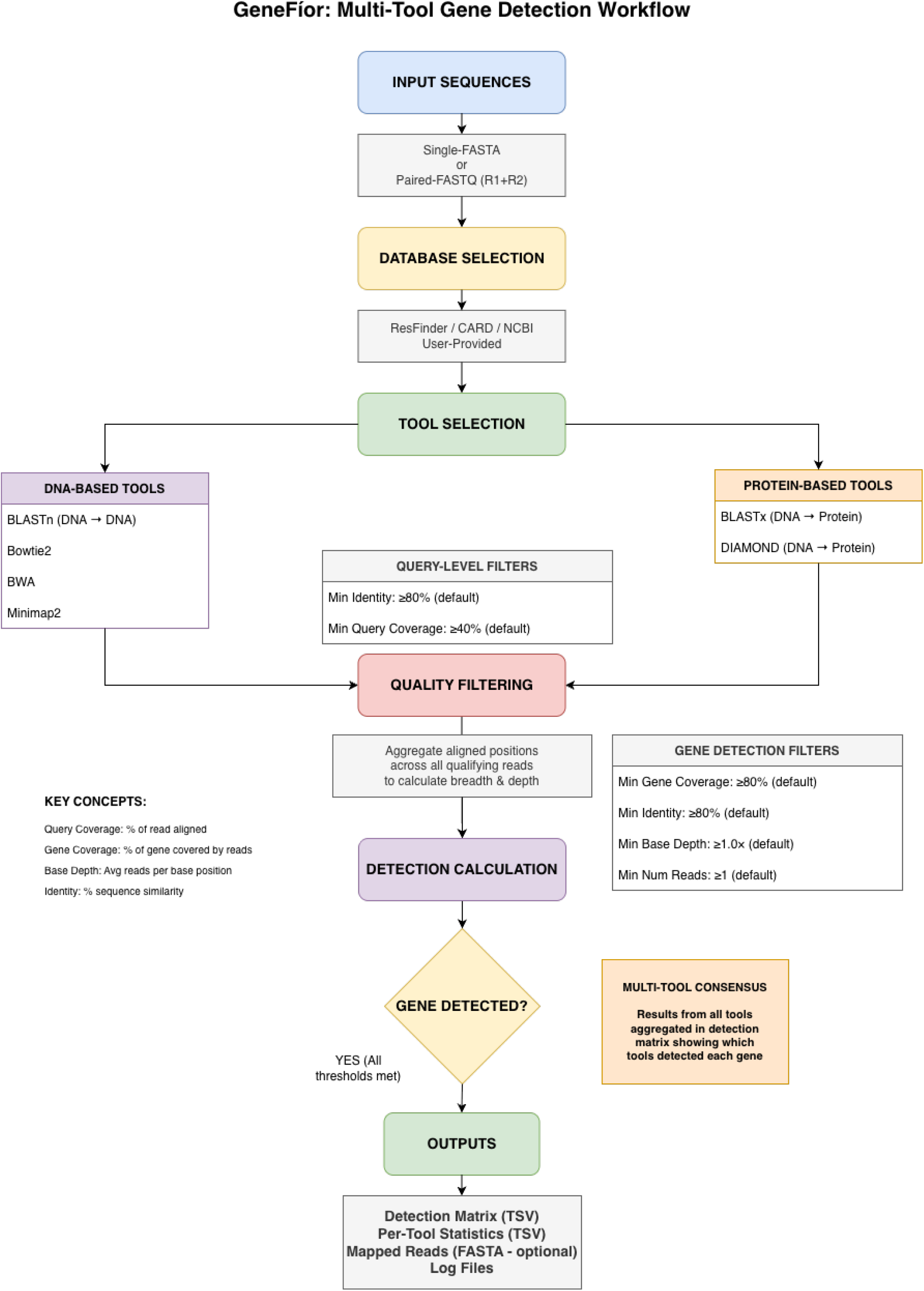
A high-level overview of the GeneFíor multi-tool detection workflow.

### 2.2 Two-Stage Threshold Architecture and Unified Detection Metrics

GeneFíor applies a two-stage threshold system to all alignment tools, separating the criteria used to pre-filter raw alignment output from those used to make final gene detection calls. This distinction is important both for computational efficiency and for ensuring that detection decisions are made on the basis of gene-level evidence rather than individual alignment quality alone.

#### Stage 1 — Query-level pre-filtering thresholds (query-min-identity, query-min-coverage)

These thresholds are applied as early as possible in the pipeline to discard low-quality alignments that cannot contribute meaningful evidence to gene detection. By default, query-min-identity is set to 80% and query-min-coverage to 40%. The asymmetry between these two defaults is deliberate: identity is held to a high standard from the outset as to allow fine-tuned filtering of spurious hits, while a lower query coverage threshold at this stage allows reads that partially overlap a gene to contribute to its cumulative gene-level coverage, which is assessed in Stage 2. Both parameters are user-definable and do influence final gene detection outcomes.

#### Stage 2 — Gene-level detection thresholds (detection-min-identity, detection-min-coverage)

These thresholds are applied during the parsing stage after all reads passing Stage 1 have been accumulated against each reference gene. By default, both are set to 80%. A gene is called as detected only if the union of all Stage-1-passing read alignments covers at least 80% of the reference gene, the mean identity across those alignments meets the detection identity threshold, the covered region achieves a minimum base depth of 1× (noted as nominal), and at least one qualifying read has been assigned (to ensure short genes can be marked as detected in edge-cases). All four criteria must be satisfied simultaneously, and all are user-configurable. Genes with at least one Stage 1 passing alignment criteria are carried forward to the consensus detection matrix described in Section 2.4.

### 2.3 Unified Coverage and Identity Calculation

To ensure that alignment quality is assessed consistently across all integrated search tools, GeneFíor applies a common framework for computing query identity and coverage, and gene identity and coverage regardless of the underlying alignment algorithm or output format. While the precise method of calculation necessarily differs between tabular output (BLAST and DIAMOND) and alignment-based output (Bowtie2, BWA, and Minimap2), the definitions are harmonised so that threshold comparisons are meaningful across tools.

Percentage identity is derived from the reported alignment for each read. For BLAST and DIAMOND, it is taken directly from the pident field of the tabular output, which follows the standard BLAST definition for each High-scoring Segment Pair (HSP). For BAM-based tools, where identity must be computed from the raw alignment, the CIGAR string is parsed position by position and identity is derived using the ‘same’ underlying definition. The total alignment length is the sum of all matched positions (M, =, X) and all inserted and deleted bases (I, D). Soft-clipped, hard-clipped, and skipped regions (S, H, N) are excluded. Identical positions are then derived by subtracting the edit distance, encoded in the NM tag and representing the total count of mismatches and indels, from this alignment length:

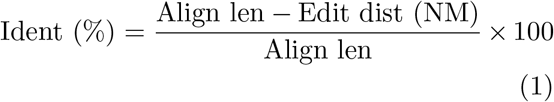

where:

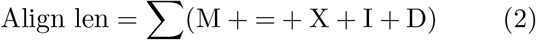

Query coverage measures the proportion of a read’s bases (or amino acids) that are genuinely aligned to the reference gene, and serves as a per-read filter to discard reads that only marginally overlap the target. For BAM-based tools, this is computed precisely at the base level: only CIGAR operations that simultaneously consume both the query and reference sequences — M, =, and X — are counted as query-aligned bases. Insertions are excluded because, although they consume query bases, those bases have no corresponding position on the reference and do not represent genuine overlap. Deletions are excluded because they consume reference positions but correspond to no bases in the read at all (there is simply nothing in the read to count). Soft and hard clips are excluded as they fall outside the alignment entirely:

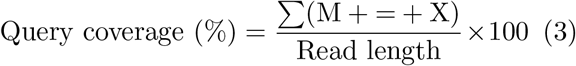

For BLAST and DIAMOND, CIGAR strings are not available in tabular output and query coverage is instead approximated from the reported alignment coordinates as the span of the aligned region relative to the total query length:

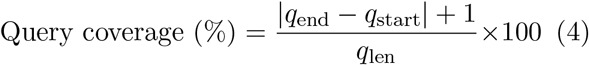

This span-based approximation is consistent with the definition used internally by BLAST and DIAMOND.It may marginally overestimate true query coverage in the presence of internal gaps, but this is not expected to materially affect detection outcomes at the thresholds used.

For gene coverage, each passing reads set of unique reference positions covered by its M, =, and X operations is recorded; deletion positions are explicitly excluded, as a deletion in the CIGAR string means those reference bases are absent from the read and cannot be considered covered. For BLAST and DIAMOND, equivalent position sets are derived from the reported subject start and end coordinates. Across all passing reads, the union of all covered reference positions is computed and expressed as a proportion of the total reference gene length.

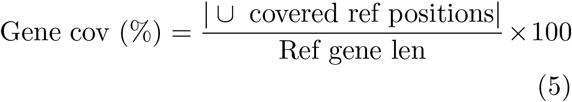

A gene is called as detected only if this accumulated gene coverage meets or exceeds the minimum gene coverage threshold, the covered positions meet a minimum base depth, and a minimum number of passing reads have contributed. These criteria are all user-configurable.

Gene identity is computed as the mean percentage identity across all reads that have passed the per-read Stage 1 filters for a given gene. Each passing read contributes its individual identity value, derived from the CIGAR string and NM tag for BAM-based tools, or taken directly from the pident field for BLAST and DIAMOND, to a running list that is averaged during finalisation:

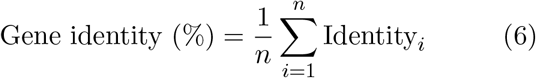

where *n* is the total number of passing reads contributing alignments to that gene. This mean is computed across all reads meeting the Stage 1 thresholds regardless of whether the gene ultimately passes detection, providing a summary measure of overall alignment quality for reporting purposes. The mean gene identity is recorded in the per-tool statistics output alongside coverage metrics, allowing users to inspect the quality of evidence supporting each gene call independently of the binary detection decision.

See Table 1 for an example tool-stats report generated for BLASTn.

**Table 1:** Table reporting example mapping results for BLASTx with the total number and those passing filtering thresholds as well as calculated gene coverage and identity metrics.

| Gene | Gene_Length | Num_Sequences_Mapped | Num_Sequences_Passing_Thresholds | Gene_Coverage | Avg_Identity | Detected |
| --- | --- | --- | --- | --- | --- | --- |
| AAC(6')-Ib7—ARO:3002578 | 786 | 1 | 1 | 17.43 | 99.27 | 0 |
| ACI-1—ARO:3004359 | 855 | 8 | 8 | 77.54 | 100 | 0 |
| ANT(6)-Ia—ARO:3002626 | 864 | 52 | 50 | 67.94 | 98.43 | 0 |
| APH(3'')-Ib—ARO:3002639 | 804 | 5 | 5 | 57.59 | 99.74 | 0 |
| APH(3')-IIIa—ARO:3002647 | 795 | 155 | 135 | 100 | 99.87 | 1 |
| APH(3')-Ia—ARO:3002641 | 816 | 4 | 3 | 55.51 | 99.34 | 0 |
| APH(6)-Id—ARO:3002660 | 837 | 1 | 1 | 18.04 | 99.34 | 0 |
| AcrF—ARO:3000502 | 3105 | 4 | 4 | 13.53 | 100 | 0 |
| AcrS—ARO:3000656 | 663 | 1 | 1 | 22.78 | 100 | 0 |
| Bado_rpoB-RIF—ARO:3004480 | 3561 | 17 | 16 | 14.1 | 88.34 | 0 |

### 2.4 Consensus Detection Matrix

To facilitate cross-tool comparison and multievidence gene detection, GeneFíor produces a consensus detection matrix summarising the detection status of every reference gene across all integrated alignment tools. Each entry in the matrix records whether a given gene was detected by a given tool, based on the coverage validation criteria described in Section 2.3, alongside the associated per-tool alignment metrics. This matrix serves two purposes. First, it exposes how tool choice, database selection, and parameter configuration influence the reported gene profile, enabling researchers to directly interrogate sources of inter-tool discordance rather than accepting a single summarised output uncritically. Second, it provides a principled basis for confidence stratification: genes detected independently by three or more tools, bridging both nucleotide and amino acid search space, can be classified as high-confidence detections, while those identified by only a single tool may warrant additional investigation. The precise concordance threshold used to define high-confidence calls can be adjusted by the user depending on the analytical context. All per-tool statistics files (See Table 1 and the full detection matrix (See Table 2 are retained as outputs, ensuring that the complete analytical record is available for inspection, reanalysis, or reporting.

**Table 2:** Summary of Gene Detections Across Different Search Search Tools.

| Gene | BLAST-DNA | BLAST-PROTEIN | BWA | Bowtie2 | DIAMOND | Minimap2 | Total Detections |
| --- | --- | --- | --- | --- | --- | --- | --- |
| aadE-Cc_1.CP013733 | 1 | 1 | 1 | 1 | 1 | 1 | 6 |
| ant(6)-Ia_1.AF330699 | 1 | 1 | 1 | 1 | 1 | 1 | 6 |
| ant(6)-Ia_2.KF421157 | 0 | 1 | 0 | 0 | 1 | 0 | 2 |
| ant(6)-Ia_3.KF864551 | 1 | 1 | 1 | 1 | 1 | 1 | 6 |
| ant(6)-Ia_5.AB247327 | 1 | 1 | 1 | 1 | 1 | 1 | 6 |
| aph(3')-IIIa_1.AF330699 | 1 | 1 | 1 | 1 | 1 | 1 | 6 |
| aph(3')-IIIa_3.AB247327 | 1 | 1 | 1 | 0 | 1 | 1 | 5 |
| blaOXA-85_1.JANA01000064 | 1 | 1 | 1 | 1 | 1 | 1 | 6 |
| cfr(C)_1.KX686749 | 1 | 1 | 1 | 0 | 1 | 0 | 4 |
| cfr(C)_2.CANB01000378 | 1 | 1 | 0 | 0 | 1 | 1 | 4 |
| tet(W/32/O)_2.AM710602 | 1 | 0 | 0 | 0 | 0 | 0 | 0 |

### 2.5 Computational Optimisations for Large-Scale Metagenomic Datasets

Running large metagenomic datasets against multiple sequence search tools presents significant computational and I/O challenges, particularly for BLAST-based tools, which can generate very large intermediate output files and are sensitive to query input size. GeneFíor implements several complementary optimisations to address these challenges, reducing peak disk usage, improving throughput, and making the workflow tractable for datasets that would otherwise exceed practical resource constraints.

To address the input requirements of BLAST and DIAMOND, GeneFíor automates the conversion of paired-end FASTQ files into a single, compressed FASTA format. During this conversion, the pipeline identifies and resolves potential read identifier collisions by appending /1 and /2 suffixes to R1 and R2 headers, ensuring read provenance is preserved throughout downstream analysis. To maximise efficiency, this process utilises high-performance streaming via seqtk [23] and pigz [24] and implements a metadata-based caching system to bypass redundant conversions on subsequent runs. For environments with high-latency storage, GeneFíor further optimises I/O by staging these processed files in a temporary directory for faster access during alignment.

For input FASTA files exceeding a configurable size threshold (default 200 MiB, adjustable via –max-fasta-chunk-mb), GeneFíor automatically splits the input into smaller chunk files and runs BLASTn/x independently on each chunk. Chunking is performed using a generator that yields chunk files as they are written to disk, allowing BLAST jobs to be submitted to a thread pool immediately without requiring all chunks to exist before any processing begins. This minimises both peak disk usage for chunk storage and the latency between chunking and job submission. After all chunk jobs complete, their filtered output files are concatenated into a single result file and temporary chunk and part-output files are removed, unless the user requests their retention via –preservechunks for debugging. The chunk size threshold and concurrency can be tuned to match available CPU, memory and I/O characteristics of the system in use. This allows GeneFíor to exploit parallelism on multi-core systems with little user-intervention.

While read-mapping tools such as Bowtie2, bwa and minimap2 can have their output streamed directly into highly-compression .bam files, BLASTn/x produces extremely large tabular output files. Rather than writing BLAST output to disk in full before filtering, resulting in extremely large filesizes, GeneFíor directs both BLASTn and BLASTx to write tabular results to stdout (-out -) and intercepts this stream in Python as the search runs. Each output line is evaluated against the Stage 1 query identity and query coverage thresholds as it arrives, and only passing lines (hits) are written to the final output file. This means the full unfiltered BLAST output is never materialised on disk. For large metagenomic datasets where unfiltered BLAST output can reach hundreds of gigabytes, this represents a substantial reduction in intermediate storage requirements.

### 2.6 User-Defined Database Generation

GeneFíor is database-agnostic by design and can be used to search sequence reads against any user-supplied set of nucleotide or amino acid reference sequences in FASTA format. As each sequence-search tool requires its own specific database index, GeneFíor automates this preparation step, generating the appropriate index formats from a single input: BLAST databases are created using makeblastdb (with nucleotide and protein options applied accordingly), DIAMOND databases using diamond makedb on protein sequences, Bowtie2 indices using bowtie2-build, minimap2 indices are created with minimap -d, and BWA indices using bwa index. Gene identifiers are harmonised across database formats to ensure consistent naming between nucleotide and amino acid search outputs, which is required for accurate cross-tool comparison and downstream reporting. As part of the AMRfíor subtool, the ResFinder and CARD databases are pre-generated and come pre-loaded as part of the Bioconda package.

### 2.7 GeneFíor Recompute

A fundamental design objective of GeneFíor is to make detection decisions fully revisable without repeating computationally expensive alignment steps. To this end, GeneFíor includes a dedicated submodule, GeneFíor-Recompute, which allows users to reapply any combination of detection thresholds to the raw alignment outputs produced by a previous GeneFíor run, without re-executing any of the underlying search tools. GeneFíor-Recompute takes as input the output directory of an existing GeneFíor run and automatically discovers all alignment output files present in the raw outputs subdirectory, identifying both tabular outputs (TSV files from BLASTn, BLASTx, and DIAMOND) and sorted BAM files (from Bowtie2, BWA, and Minimap2). The tool identifies which databases and alignment tools are represented in the existing outputs and processes only those tools for which raw output files are found, unless the user further restricts analysis to a specific subset. Database and tool identity are inferred automatically from output filenames, requiring no additional configuration from the user. Once files are discovered, GeneFíor-Recompute instantiates the same parsing and detection logic used by the main GeneFíor workflow, parse blast results for tabular outputs and parse bam results for BAM files, applying whichever threshold values the user specifies at the command line. All four detection parameters are independently configurable: minimum gene coverage (–d-min-cov), minimum identity (–d-min-id), minimum base depth (–d-min-base-depth), and minimum number of contributing reads (–d-min-reads). The query-level pre-filter for minimum query identity (–q-min-id) and minimum query coverage (–q-min-cov) are also user-configurable and are applied during parsing in the same way as in the primary workflow. Recomputed per-tool statistics files and cross-tool detection matrices are written to a user-specified output directory, leaving the original results untouched. This design makes GeneFíor-Recompute particularly well-suited to threshold sensitivity analyses, where the same alignment data must be interrogated under multiple detection criteria. Rather than storing multiple complete copies of detection results, users can archive the raw alignment outputs once at lower thresholds and regenerate detection calls under any parameter combination on demand. The full detection pipeline, from per-read quality assessment through to gene-level calling and consensus matrix generation, is executed identically to the primary workflow, ensuring that recomputed results are directly comparable to primary results produced under the same thresholds.

### 2.8 GeneFíor Gene Stats

GeneFíor-Gene-Stats is a complementary submodule designed for targeted, position-level investigation of genes of interest identified by the main GeneFíor workflow. Users supply one or more gene names, either as a comma-separated list or as a file with one name per line, along with the GeneFíor output directory, the databases to interrogate, and the tools to include. For BAM-based tools, the CIGAR string of each alignment is parsed at base resolution. This produces a per-position pileup for each gene, tracking the identity and count of every base observed across all mapped reads. For BLAST and DIAMOND tabular outputs, base-level data is not directly available from the alignment format; however, if the user provides the original query FASTA file, GeneFíor-Gene-Stats extracts the aligned portion of each query read from the stored sequences and maps these bases to the corresponding subject positions, enabling approximate base-level coverage even from tabular output. From the accumulated per-position data, GeneFíor-Gene-Stats computes a suite of coverage statistics for each gene and tool combination: total and percentage of positions covered, total read count, mean and maximum depth, and a list of uncovered gap regions with their coordinates and lengths.Sequence variants are identified at positions with a minimum depth of 3×, where the consensus observed base differs from the provided reference sequence; each variant is reported with its position, reference base, consensus alternative base, sequencing depth, and variant frequency. As shown in Figure 2, coverage across the gene length is additionally visualised as an ASCII depth histogram where average depth per bin is displayed as a normalised bar chart, providing an immediate visual summary of the distribution of read coverage across the gene.

**Figure 2:**
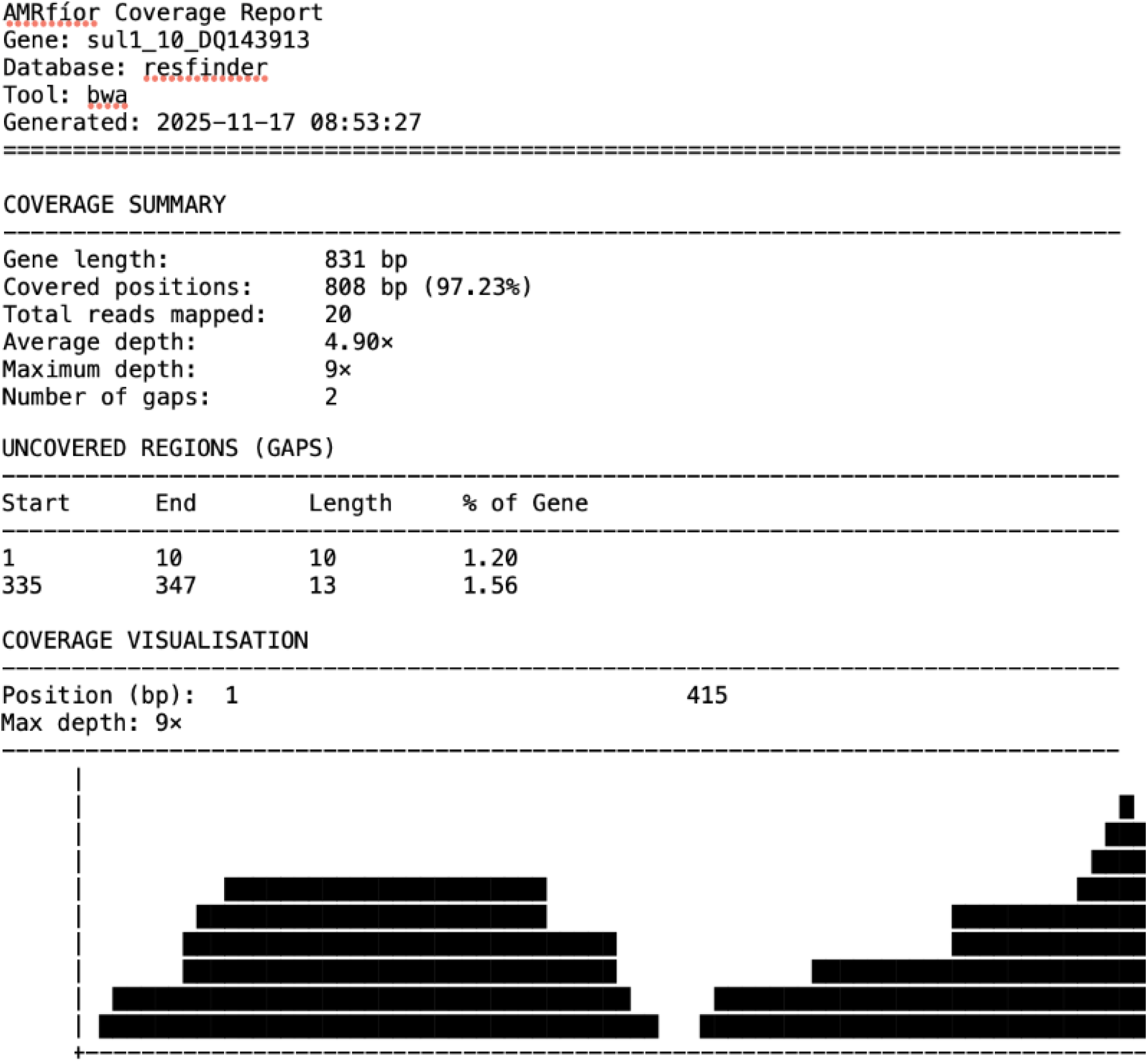
An example Gene-Stats coverage report for the gene sul1 10 DQ143913 from the ResFinder database as found by bwa. This simple report clearly shows the two regions of the gene where no reads where aligned, hinting at sequence divergence at those positions between the sample and database version of this gene.

For each gene, GeneFíor-Gene-Stats also generates a cross-tool comparison report (see Figure 3, tabulating coverage percentage, average depth, read count, number of gap regions, and number of detected variants side by side across all tools that produced alignments to that gene. A tool agreement analysis is included, reporting the number and percentage of gene positions covered by all tools simultaneously, by at least one tool, and exclusively by individual tools. This exposes positions where tools agree on coverage, providing higher-confidence evidence, as well as tool-specific regions that may reflect algorithm-level differences in alignment behaviour or sensitivity.

**Figure 3:**
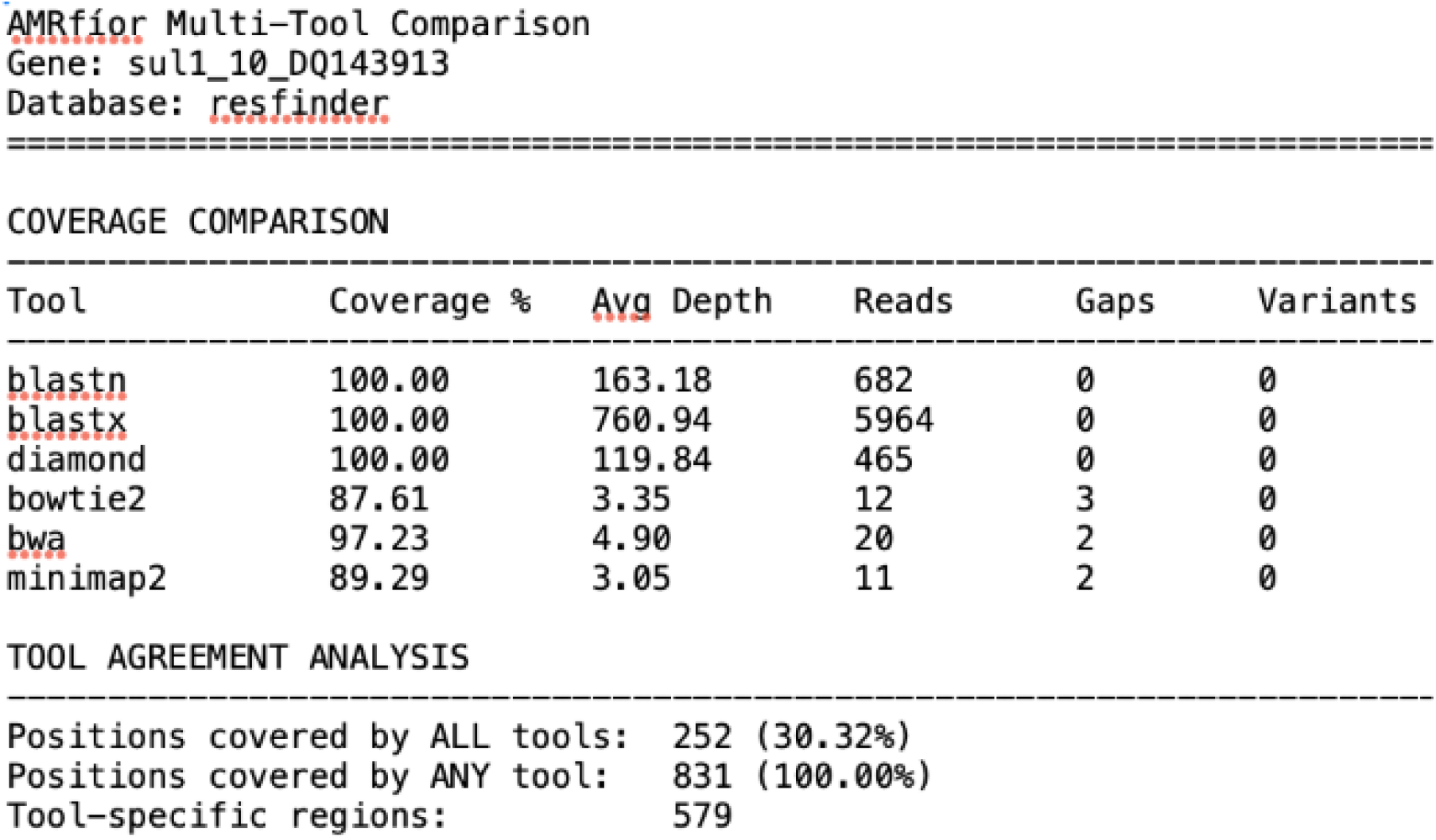
An example Gene-Stats multi-tool report for the gene sul1 10 DQ143913 from the ResFinder database. This simple table reports the stark differences in search results from the 6 tools.

### 2.9 Gene Detection and Database Preparation with AMRFíor

Antimicrobial resistance gene detection was performed using AMRFíor, a sub-workflow of GeneFíor that integrates the 6 sequence search and alignment algorithms to identify ARGs from pre-generated databases. ARG reference databases were obtained from ResFinder (version 2.6.0, accessed 2025-10-10) and the Comprehensive Antibiotic Resistance Database (CARD, version 4.0.1, accessed 2025-10-10). The ResFinder database provides only DNA sequences for its ARGs so the full set were translated to their protein counterpart using the ‘universal’ codon table 11. CARD provided its ARGs as both DNA and protein sequences but harmonisation of gene names was needed to ensure uniformity between DNA and AA search tools. Each database was formatted for all alignment tools according to their respective requirements.

### 2.10 Case studies, example usage

### 2.11 Use Case: ARG detection in genomes

Bacterial genomes were sourced from the abritAMR dataset [25], which included 410 bacterial isolates (22 species) previously tested by a combination of PCR assays, including a carbapenemase and ESBL real-time multiplex PCR. PCR-positive results reported by Sherry *et al* [25], were extracted from the accompanying source data file. The PCR-confirmed resistance gene group names were then used to match against ARGs from the ResFinder and CARD databases by gene name only, enabling the comparison of sequence-based detection against a PCR-confirmed reference standard.

The ARG classes included in this comparison were: *CTX-M* group 1, *CTX-M* group 9, *IMI, IMP, KPC, NDM, OXA*-23-like, *OXA*-48-like, *OXA-51* -like, *OXA-58* -like, and *VIM*.

AMRFíor was ran using all 6 tools.

## 3 Results

### 3.1 Use Case: ARG detection in genomes

To evaluate the ARG identification performance of the AMRFíor module, we compared our results against validated PCR data from Sherry *et al* [25]. A total of 11 resistance gene groups were evaluated across 410 PCR-confirmed genomes. GeneFíor analysis, incorporating all six detection tools, had a mean runtime of 2 hours 23 minutes per genome (ranging from 23 minutes to 5 hours 33 minutes) against the CARD database and 2 hours 26 minutes (17 minutes to 12 hours 17 minutes) against the ResFinder database, with 20 CPU threads. Genomes had a mean of 4.08 million paired-end reads (1,529,144 to 10,149,360).

One hundred percent of PCR-confirmed ARGs were detected for all gene groups across both databases and all genomes (Table 3). However, while BLAST-based tools consistently achieved 100% sensitivity across all gene groups and both databases, substantial differences in reported genes were observed between the remaining search tools. DIAMOND performed strongly across most gene groups, though minor omissions were observed for specific genes such as *‘CTX-M* group 1’ and *‘OXA-51* -like’ in the ResFinder and CARD databases, respectively. The greatest variation was observed among alignment-based tools, with BWA, Bowtie2, and Minimap2 showing markedly reduced sensitivity for certain gene groups. *OXA51* -like genes were particularly poorly detected by these tools across both databases, with sensitivities as low as 0–4.2%. Overall, while these preliminary results are not yet conclusive, they do highlight the importance of tool choice, sequence search space (DNA/AA) and database choice. As we can see in Table 3, the read-mapping approaches (BWA, Bowtie2, Minimap2) report no genes of *‘CTX-M* group 1’ when using the ResFinder database, but they do detect almost all of them when using the CARD database. Frustratingly, this does not suggest that CARD is the more representative database as for the gene *‘KPC’*, while all 6 tools reported all 10 genes using the ResFinder database, Bowtie2 and Minimap2 reported 8 and 9, respectively for the CARD database.

**Table 3:** Number of PCR-confirmed positive genomes containing each antimicrobial resistance gene as detected by GeneFíor using ResFinder and CARD databases. The PCR column indicates the total number of genomes confirmed positive by PCR for each resistance gene group [25]. Values for each tool represent the number of these PCR-positive genomes in which the ARG was detected. Tools shown are BLASTn, BLASTx, BWA, Bowtie2, DIAMOND, and Minimap2 as implemented within the GeneFiór pipeline.

| **ResFinder-Gene** | **PCR** | **BLASTn** | **BLASTx** | **BWA** | **Bowtie2** | **DIAMOND** | **Minimap2** |
| --- | --- | --- | --- | --- | --- | --- | --- |
| **CTX-M group 1** | 58 | 58 | 58 | 0 | 0 | 45 | 0 |
| **CTX-M group 9** | 69 | 69 | 69 | 59 | 58 | 69 | 60 |
| **IMI** | 5 | 5 | 5 | 4 | 4 | 5 | 4 |
| **IMP** | 155 | 155 | 155 | 155 | 155 | 155 | 155 |
| **KPC** | 10 | 10 | 10 | 10 | 10 | 10 | 10 |
| **NDM** | 38 | 38 | 38 | 38 | 38 | 38 | 38 |
| **OXA-23-like** | 22 | 22 | 22 | 22 | 22 | 22 | 21 |
| **OXA-48-like** | 21 | 21 | 21 | 5 | 5 | 21 | 5 |
| **OXA-51-like** | 24 | 24 | 24 | 0 | 0 | 0 | 0 |
| **OXA-58-like** | 6 | 6 | 6 | 6 | 6 | 6 | 6 |
| **VIM** | 2 | 2 | 2 | 2 | 2 | 2 | 2 |
| **CARD-Gene** | **PCR** | **BLASTn** | **BLASTx** | **BWA** | **Bowtie2** | **DIAMOND** | **Minimap2** |
| **CTX-M group 1** | 58 | 58 | 58 | 58 | 55 | 58 | 56 |
| **CTX-M group 9** | 69 | 69 | 69 | 69 | 69 | 69 | 68 |
| **IMI** | 5 | 5 | 5 | 4 | 4 | 5 | 4 |
| **IMP** | 155 | 155 | 155 | 155 | 155 | 155 | 155 |
| **KPC** | 10 | 10 | 10 | 10 | 8 | 10 | 9 |
| **NDM** | 38 | 38 | 38 | 38 | 38 | 38 | 38 |
| **OXA-23-like** | 22 | 22 | 22 | 21 | 21 | 22 | 19 |
| **OXA-48-like** | 21 | 21 | 21 | 20 | 20 | 21 | 20 |
| **OXA-51-like** | 24 | 24 | 24 | 1 | 1 | 7 | 1 |
| **OXA-58-like** | 6 | 6 | 6 | 6 | 6 | 6 | 6 |
| **VIM** | 2 | 2 | 2 | 2 | 2 | 2 | 2 |

Together, these findings reinforce the core motivation for a multi-tool approach: any single tool may produce a measurably incomplete or distorted resistance profile depending on the gene group and database in question.

## 4 Discussion

Through GeneFíor, we have demonstrated that tool choice, search space (nucleotide versus amino acid), database selection, and parameter configuration are not neutral decisions as each measurably and sometimes substantially alters the set of genes reported as detected. This is only visible because multiple tools are run simultaneously: a single-tool analysis would present any one of these outcomes as the result, with no means of knowing how much it diverges from what other equally defensible methodological choices would have produced.

Although this work has so far focused on ARG detection, the challenges identified are not domain-specific. Any study relying on sequence similarity to detect genes of interest is subject to the same limitations, including database redundancy, arbitrary clustering thresholds, and hidden parameterisation. The increasing adoption of machine learning-based approaches further amplifies these concerns, as gains in sensitivity are often accompanied by reduced transparency and limited interpretability.

Rather than proposing yet another detection algorithm, GeneFíor advocates a return to first prin-ciples: combining established methods within a framework that prioritises transparency, flexibility, and reproducibility. In doing so, it provides a foundation for more reliable gene detection across diverse applications, while enabling critical evaluation of both tool behaviour and biological inference. Future work should focus not only on improving detection performance, but on standardising reporting, exposing methodological assumptions, and ensuring that gene detection remains both interpretable and verifiable as datasets and analytical complexity continue to grow.

## 5 Data availability

Not applicable.

## 7 Author contributions statement

N.J.D designed and developed GeneFíor, K.L carried out the case study analysis and performed extensive user testing. N.J.D and K.L contributed to the vision of Fíor and writing of the manuscript equally.

## References

[1] The Global AMR Innovation Fund. https://www.gov.uk/government/groups/the-global-amr-innovation-fund, 2020.

[2] Directorate-General for Research and Innovation. https://research-and-innovation.ec.europa.eu/news/all-research-and-innovation-news/new-european-partnership-one-health-amr-eu253-million-research-and-innovation-against-antimicrobial-2025-09-23_en, 2025.

[3] Jenna Rychert. Benefits and limitations of maldi-tof mass spectrometry for the identification of microorganisms. Journal of Infectiology and Epidemiology, 2(4), 2019.

[4] M Benkova, O Soukup, and J Marek. Antimicrobial susceptibility testing: currently used methods and devices and the near future in clinical practice. Journal of applied microbiology, 129(4):806–822, 2020.

[5] European Committee on Antimicrobial Susceptibility Testing Clinical Breakpoints-Breakpoints and Guidance. https://www.eucast.org/clinical_breakpoints, 2025.

[6] Gary V Doern and Stephen M Brecher. The clinical predictive value (or lack thereof) of the results of in vitro antimicrobial susceptibility tests. Journal of Clinical Microbiology, 49(9 Supplement):S11–S14, 2011.

[7] Gustavo Arango-Argoty, Emily Garner, Amy Pruden, Lenwood S Heath, Peter Vikesland, and Liqing Zhang. Deeparg: a deep learning approach for predicting antibiotic resistance genes from metagenomic data. Microbiome, 6(1):23, 2018.

[8] Alfred Ferrer Florensa, Rolf Sommer Kaas, Philip Thomas Lanken Conradsen Clausen, Derya Aytan-Aktug, and Frank M Aarestrup. Resfinder–an open online resource for identification of antimicrobial resistance genes in next-generation sequencing data and prediction of phenotypes from genotypes. Microbial genomics, 8(1):000748, 2022.

[9] Brian P Alcock, William Huynh, Romeo Chalil, Keaton W Smith, Amogelang R Raphenya, Mateusz A Wlodarski, Arman Edalatmand, Aaron Petkau, Sohaib A Syed, Kara K Tsang, et al. Card 2023: expanded curation, support for machine learning, and resistome prediction at the comprehensive antibiotic resistance database. Nucleic acids research, 51(D1):D690–D699, 2023.

[10] Lucy Dillon, Nicholas J Dimonaco, and Christopher J Creevey. Accessory genes define species-specific routes to antibiotic resistance. Life science alliance, 7(4), 2024.

[11] Michael Feldgarden, Vyacheslav Brover, Narjol Gonzalez-Escalona, Jonathan G Frye, Julie Haendiges, Daniel H Haft, Maria Hoffmann, James B Pettengill, Arjun B Prasad, Glenn E Tillman, et al. Amrfinderplus and the reference gene catalog facilitate examination of the genomic links among antimicrobial resistance, stress response, and virulence. Scientific reports, 11(1):12728, 2021.

[12] Ea Zankari, Rosa Allesøe, Katrine G Joensen, Lina M Cavaco, Ole Lund, and Frank M Aarestrup. Pointfinder: a novel web tool for wgs-based detection of antimicrobial resistance associated with chromosomal point mutations in bacterial pathogens. Journal of antimicrobial chemotherapy, 72(10):2764– 2768, 2017.

[13] Will PM Rowe and Martyn D Winn. Indexed variation graphs for efficient and accurate resistome profiling. Bioinformatics, 34(21):3601–3608, 2018.

[14] Stephen F Altschul, Warren Gish, Webb Miller, Eugene W Myers, and David J Lipman. Basic local alignment search tool. Journal of molecular biology, 215(3):403–410, 1990.

[15] Benjamin Buchfink, Chao Xie, and Daniel H Huson. Fast and sensitive protein alignment using diamond. Nature methods, 12(1):59–60, 2015.

[16] Ben Langmead and Steven L Salzberg. Fast gapped-read alignment with bowtie 2. Nature methods, 9(4):357–359, 2012.

[17] Heng Li and Richard Durbin. Fast and accurate short read alignment with burrows–wheeler transform. bioinformatics, 25(14):1754–1760, 2009.

[18] Heng Li. Minimap2: pairwise alignment for nucleotide sequences. Bioinformatics, 34(18):3094–3100, 2018.

[19] Fernando Baquero, Jose L Martinez, V F. Lanza, Jerónimo Rodríguez-Beltrán, Juan Carlos Galán, Alvaro San Millán, Rafael Cantón, and Teresa M Coque. Evolutionary pathways and trajectories in antibiotic resistance. Clinical Microbiology Reviews, 34(4):e00050–19, 2021.

[20] Aaron Hinz, André Amado Rees Kassen, Claudia Bank, and Alex Wong. Unpredictability of the fitness effects of antimicrobial resistance mutations across environments in escherichia coli. Molecular Biology and Evolution, 41(5):msae086, 2024.

[21] Jorge Rojas-Vargas, Hugo G Castelán-Sánchez, and Liliana Pardo-López. Hadeg: a curated hydrocarbon aerobic degradation enzymes and genes database. Computational Biology and Chemistry, 107:107966, 2023.

[22] Siyu Zhou, Bo Liu, Dandan Zheng, Lihong Chen, and Jian Yang. Vfdb 2025: an integrated resource for exploring anti-virulence compounds. Nucleic acids research, 53(D1):D871–D877, 2025.

[23] Wei Shen, Shuai Le, Yan Li, and Fuquan Hu. Seqkit: a cross-platform and ultrafast toolkit for fasta/q file manipulation. PloS one, 11(10):e0163962, 2016.

[24] multi-core machines pigz: A parallel implementation of gzip for modern multi processor. https://zlib.net/pigz/, 2025.

[25] Norelle L. Sherry, Kristy A. Horan, Susan A. Ballard, Anders Gonçalves da Silva, Claire L. Gorrie, Mark B. Schultz, Kerrie Stevens, Mary Valcanis, Michelle L. Sait, Timothy P. Stinear, et al. An ISO-certified genomics workflow for identification and surveillance of antimicrobial resistance. Nature Communications, 14(1):60, 2023.

